# Functional diversity plays a role in driving β-diversity: Or does it?

**DOI:** 10.1101/594044

**Authors:** David Murray-Stoker

## Abstract

Patrick and Brown (2018) suggest that functional diversity of the species pool has an important role in generating β-diversity. Using a combination of path analysis and model selection, they ostensibly provide support for this hypothesis; however, they neglected to put theory and modeling into proper ecological and statistical context. Here, I present a re-analysis of their data. I conclude that the drivers of β-diversity are variable, with functional diversity typically having a reduced, if any, role compared to consistently stronger roles played by γ-diversity or environmental variation on structuring β-diversity.

## Introduction

Environmental filtering is a common mechanism used to explain community assembly (Poff 1997, Leibold et al. 2004). Under this framework, realized assemblages are derived from the regional pool by a series of environmental variables selecting for taxa able to establish under the local conditions. Derived from this framework is the hypothesized positive relationship between environmental heterogeneity and β-diversity. Considering each community is comprised of taxa able to occupy and persist in the local environmental conditions, greater variation in environmental conditions in the region results in greater variation among communities due to differential composition among habitats (Logue et al. 2011, Heino et al. 2013). This hypothesis has been evaluated in terrestrial (Kraft et al. 2011, Cramer and Verboom 2017) and aquatic (Grönroos et al. 2013, Heino et al. 2013, Astorga et al. 2014, Heino et al. 2017, Wojciechowski et al. 2017) systems, but there is no conclusive support for the predominant drivers of β-diversity.

Patrick and Brown (2018) suggested that functional diversity could play an important role in generating β-diversity. Derived from the environmental filtering framework, they proposed that greater functional differences among taxa in the region would allow for differential composition among local communities, and this relationship would be greater at higher levels of environmental heterogeneity or filtering. In other words, Patrick and Brown (2018) hypothesized a positive relationship between functional diversity and β-diversity. This hypothesis is intuitive in the broader context of the environmental filtering framework, and, through the use of statistical modeling and causal inference, Patrick and Brown (2018) report functional diversity having an important role in predicting β-diversity. Although the relative importance of functional diversity as a predictor was generally less than environmental heterogeneity and greater than γ-richness, they asserted that functional diversity was still important and useful as a predictor of β-diversity. I contend that the study by Patrick and Brown (2018) does not adequately support this conclusion.

I contest functional diversity as a driver of β-diversity and rather as a correlate of β-diversity. This argument is based on the hierarchical and correlative nature of α-, β-, and γ- diversity (Whittaker 1960) and the premise that functional diversity is dynamic, simultaneously influencing and being influenced by (i.e. correlated to) α-, β-, and γ-diversity. I assert that functional diversity is a constituent of community diversity and not an independent entity. I hypothesized that models with correlations rather than causal pathways between or among diversity metrics (i.e. functional, β-, and γ-diversity) would have better model support than models proposed by Patrick and Brown (2018). Additionally, the relative importance of functional diversity would be consistently less than the effects of environmental heterogeneity and γ-diversity on β-diversity.

## Re-Analyses

I first re-analyzed the data by testing the path models hypothesized by Patrick and Brown (2018, see fig. 2) and the alternative path models presented here (fig. 1) by using structural equation models and model selection techniques. I generally followed the methods outlined by Patrick and Brown (2018), although I have provided more detailed methods in the appendix. Models were fit for the whole community and each individual functional feeding group. I refer to models by the number associated with the model structure. Models 1-5 were proposed by Patrick and Brown (2018), while I proposed models 6-8.

**Figure 1.**
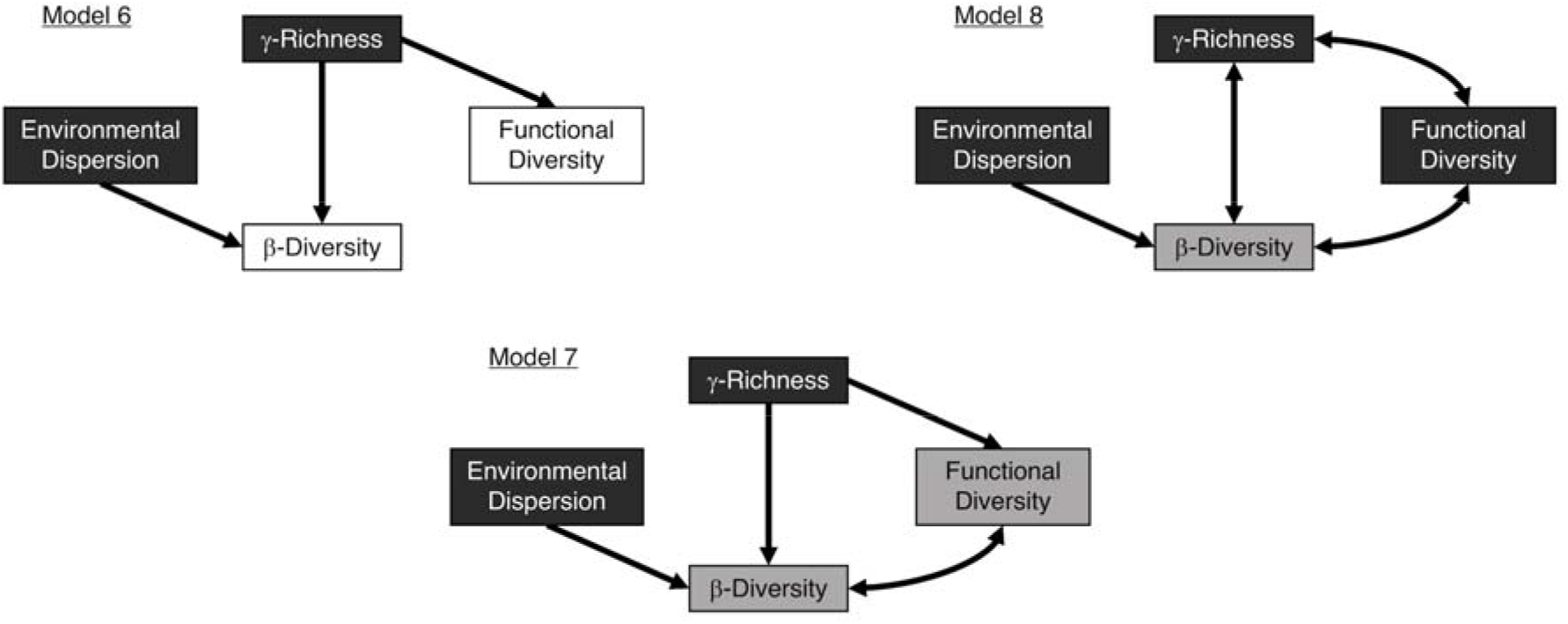
Alternative path models to those proposed by Patrick and Brown (2018). Models display the hypothesized relationships between environmental dispersion, γ-richness, functional diversity, and β-diversity. Exogenous (independent) variables are represented in black boxes, endogenous (dependent) variables in grey boxes, and response-only variables in white boxes. Single-headed arrows represent causal pathways, and double-headed arrows represent correlational pathways.

I found that the top model or models varied among the full community and functional feeding group subsets (table 1). Variation in top models notwithstanding, causal pathways linking environmental heterogeneity or γ-diversity to β-diversity consistently had the strongest effect. Environmental heterogeneity had significant effects on β-diversity for the whole community and collector-gatherer and filter-feeder subsets; γ-diversity had significant effects on β-diversity for the herbivore, shredder, and predator subsets; and functional diversity had significant effects on β-diversity for the collector-gatherer and filter-feeder subsets. Even when functional diversity effects on β-diversity were significant, these effects were considerably reduced in relative importance (collector-gatherer model 4: functional diversity = 0.264, environmental heterogeneity = 0.527) or negatively related to β-diversity (filter-feeder model 4: functional diversity = −0.509, environmental heterogeneity = 0.419; filter-feeder model 5: functional diversity = −0.502, environmental heterogeneity = 0.414). Net effects also showed that functional diversity was generally of lower or negligible relative importance compared to the net effects of environmental heterogeneity or γ-diversity (table 1).

**Table 1.** Summary of all the path models in the re-analyses.

| Group | Fit (P) | AIC | ΔAIC | AIC Weight | γ-Richness on FD | FD on $\beta$ | Environmental Dispersion on $\beta$ | γ-Richness on $\beta$ |
| --- | --- | --- | --- | --- | --- | --- | --- | --- |
| Whole Community |  |  |  |  |  |  |  |  |
| 2 | *0.468* | *510.65* | *0.00* | *0.19* | *NULL* | *NA* | *0.411* | *-0.096* |
| 4 | *0.452* | *510.75* | *0.10* | *0.18* | *NULL* | *0.085* | *0.391* | *NULL* |
| 5 | *0.700* | *510.82* | *0.17* | *0.17* | *0.210* | *0.085* | *0.390* | *0.018* |
| 3 | **0.382** | **512.03** | **1.38** | **0.09** | **NULL** | **0.108** | **0.410** | **-0.119** |
| 8 | **0.382** | **512.03** | **1.38** | **0.09** | **NULL** | **NULL** | **0.410** | **NULL** |
| 7 | **0.964** | **512.11** | **1.46** | **0.09** | **0.210** | **NULL** | **0.414** | **-0.097** |
| 6 | **0.964** | **512.11** | **1.46** | **0.09** | **0.210** | **NA** | **0.414** | **-0.097** |
| 1 | **0.964** | **512.11** | **1.46** | **0.09** | **0.210** | **0.109** | **0.414** | **-0.097** |
| Collector-Gatherer |  |  |  |  |  |  |  |  |
| 4 | *0.799* | *500.70* | *0.00* | *0.31* | *NULL* | *0.264* | *0.527* | *NULL* |
| 8 | **0.764** | **502.23** | **1.53** | **0.15** | **NULL** | **NULL** | **0.520** | **NULL** |
| 3 | **0.764** | **502.23** | **1.53** | **0.15** | **NULL** | **0.250** | **0.520** | **0.085** |
| 5 | **0.740** | **502.29** | **1.59** | **0.14** | **0.173** | **0.263** | **0.525** | **0.045** |
| 7 | 0.717 | 503.82 | 3.12 | 0.07 | 0.173 | NULL | 0.517 | 0.127 |
| 6 | 0.717 | 503.82 | 3.12 | 0.07 | 0.173 | NA | 0.517 | 0.127 |
| 1 | 0.717 | 503.82 | 3.12 | 0.07 | 0.173 | 0.248 | 0.517 | 0.127 |
| 2 | 0.211 | 504.21 | 3.51 | 0.05 | NULL | NA | 0.511 | 0.130 |
| Herbivore |  |  |  |  |  |  |  |  |
| 2 | *0.973* | *494.92* | *0.00* | *0.46* | *NULL* | *NA* | *-0.192* | *0.500* |
| 8 | 0.895 | 496.92 | 2.00 | 0.17 | NULL | NULL | -0.193 | NULL |
| 3 | 0.895 | 496.92 | 2.00 | 0.17 | NULL | 0.009 | -0.193 | 0.499 |
| 7 | 0.770 | 498.78 | 3.86 | 0.07 | 0.047 | NULL | -0.194 | 0.502 |
| 6 | 0.770 | 498.78 | 3.86 | 0.07 | 0.047 | NA | -0.194 | 0.502 |
| 1 | 0.770 | 498.78 | 3.86 | 0.07 | 0.047 | 0.009 | -0.194 | 0.502 |
| 4 | 0.004 | 508.06 | 13.13 | 0.00 | NULL | 0.031 | -0.167 | NULL |
| 5 | 0.001 | 509.92 | 15.00 | 0.00 | 0.047 | 0.031 | -0.167 | 0.001 |
| Filter-Feeder |  |  |  |  |  |  |  |  |
| 5 | *0.467* | *496.24* | *0.00* | *0.24* | *0.254* | *-0.509* | *0.419* | *-0.129* |
| 4 | *0.272* | *496.61* | *0.38* | *0.20* | *NULL* | *-0.502* | *0.414* | *NULL* |
| 1 | **0.355** | **497.57** | **1.33** | **0.12** | **0.254** | **-0.530** | **0.400** | **-0.037** |
| 7 | **0.355** | **497.57** | **1.33** | **0.12** | **0.254** | **NULL** | **0.400** | **-0.037** |
| 6 | **0.355** | **497.57** | **1.33** | **0.12** | **0.254** | **NA** | **0.400** | **-0.037** |
| 3 | **0.198** | **497.95** | **1.71** | **0.10** | **NULL** | **-0.528** | **0.399** | **0.098** |
| 8 | **0.198** | **497.95** | **1.71** | **0.10** | **NULL** | **NULL** | **0.399** | **NULL** |
| 2 | < 0.000 | 512.46 | 16.22 | 0.00 | NULL | NA | 0.337 | -0.021 |
| Shredder |  |  |  |  |  |  |  |  |
| 2 | *0.726* | *462.37* | *0.00* | *0.37* | *NULL* | *NA* | *0.021* | *0.652* |
| 3 | **0.759** | **463.61** | **1.24** | **0.20** | **NULL** | **0.104** | **0.033** | **0.635** |
| 8 | **0.759** | **463.61** | **1.24** | **0.20** | **NULL** | **NULL** | **0.033** | **NULL** |
| 7 | 0.459 | 465.60 | 3.23 | 0.07 | 0.160 | NULL | 0.033 | 0.651 |
| 6 | 0.459 | 465.60 | 3.23 | 0.07 | 0.160 | NA | 0.033 | 0.651 |
| 1 | 0.459 | 465.60 | 3.23 | 0.07 | 0.160 | 0.104 | 0.033 | 0.652 |
| 4 | < 0.000 | 483.78 | 21.41 | 0.00 | NULL | 0.207 | 0.051 | NULL |
| 5 | < 0.000 | 485.78 | 23.41 | 0.00 | 0.160 | 0.207 | 0.051 | 0.033 |
| Predator |  |  |  |  |  |  |  |  |
| 1 | *0.664* | *504.15* | *0.00* | *0.19* | *0.129* | *0.136* | *0.239* | *0.434* |
| 7 | *0.664* | *504.15* | *0.00* | *0.19* | *0.129* | *NULL* | *0.239* | *0.434* |
| 6 | *0.664* | *504.15* | *0.00* | *0.19* | *0.129* | *NA* | *0.239* | *0.434* |
| 2 | *0.228* | *504.29* | *0.14* | *0.18* | *NULL* | *NA* | *0.237* | *0.448* |
| 8 | **0.206** | **505.12** | **0.97** | **0.12** | **NULL** | **NULL** | **0.246** | **NULL** |
| 3 | **0.206** | **505.12** | **0.97** | **0.12** | **NULL** | **0.139** | **0.246** | **0.428** |
| 5 | 0.008 | 511.55 | 7.40 | 0.00 | 0.129 | 0.192 | 0.345 | 0.025 |
| 4 | 0.006 | 512.52 | 8.37 | 0.00 | NULL | 0.193 | 0.346 | NULL |

Note: All models were compared for the full community and each functional feeding group using
Akaike's information criteria (AIC). The difference in AIC scores ( ΔAIC) is relative to the
model with the lowest AIC score, and model weight is the relative support of the individual model within the model set. Models in boldface had good fit to the data ( $\chi^2 > 0.05$ ) and $\Delta AIC <$ 2.00, and italicized models had the greatest relative support. Net effects are the sum of the direct and indirect effects mediated by unidirectional causal pathways; correlational pathways between variables cannot be included when calculating net effects. If correlations were the singular or mediating pathway modeled between the variables, the effects are reported as NULL. FD = functional diversity; NA = not applicable (i.e. no causal pathway in the model).

Although the alternative models were only retained among the top models for the predator community subset (table 1), results from the re-analysis drastically contradicted what was presented by Patrick and Brown (2018, see table 2). Discrepancies were not just in relative model support, but also goodness-of-fit tests and net effects of predictor variables; therefore, I proceeded to also re-analyze the original model set (i.e. models 1-5). Again, there was variation in the top model or models for the whole community and functional feeding group subsets (table 2). Causal pathways linking environmental heterogeneity or γ-diversity to β-diversity consistently had the strongest effect. Importantly, a link between functional diversity and β-diversity was only retained in the top model(s) for three of the six community sets (table 2); this contrasts Patrick and Brown (2018), who report a link between functional diversity and β-diversity in at least one top model for all six model sets. In the remaining community subsets, models including a causal pathway between functional diversity and β-diversity were equivalent to or outperformed by models without the causal pathway. Even when retained in the top models, the causal pathway between functional diversity and β-diversity was only significant for two of the three community subsets (collector-gatherer and filter-feeder subsets), and of reduced importance for driving (i.e. collector-gatherer subset) or negatively-related to (filter-feeder subset) β-diversity (table 2) when significant.

**Table 2.** Summary of the candidate path models proposed by Patrick and Brown (2018), with models reported in order of AIC and ΔAIC.

| Group | Fit (P) | AIC | ΔAIC | AIC Weight | γ-Richness on FD | FD on $\beta$ | Environmental Dispersion on $\beta$ | γ-Richness on $\beta$ |
| --- | --- | --- | --- | --- | --- | --- | --- | --- |
| Whole Community |  |  |  |  |  |  |  |  |
| 2 | **0.468** | **510.65** | **0.00** | **0.26** | **NULL** | **NA** | **0.411** | **-0.096** |
| 4 | **0.452** | **510.75** | **0.10** | **0.25** | **NULL** | **0.085** | **0.391** | **NULL** |
| 5 | **0.700** | **510.82** | **0.17** | **0.24** | **0.210** | **0.085** | **0.390** | **0.018** |
| 3 | **0.382** | **512.03** | **1.38** | **0.13** | **NULL** | **0.108** | **0.411** | **-0.119** |
| 1 | **0.964** | **512.11** | **1.46** | **0.12** | **0.210** | **0.109** | **0.414** | **-0.097** |
| Collector-Gatherer |  |  |  |  |  |  |  |  |
| 4 | **0.799** | **500.70** | **0.00** | **0.44** | **NULL** | **0.264** | **0.527** | **NULL** |
| 3 | **0.764** | **502.23** | **1.53** | **0.20** | **NULL** | **0.250** | **0.520** | **0.085** |
| 5 | **0.740** | **502.29** | **1.59** | **0.20** | **0.173** | **0.263** | **0.525** | **0.045** |
| 1 | 0.717 | 503.82 | 3.12 | 0.09 | 0.173 | 0.248 | 0.517 | 0.127 |
| 2 | 0.211 | 504.21 | 3.51 | 0.08 | NULL | NA | 0.511 | 0.130 |
| Herbivore |  |  |  |  |  |  |  |  |
| 2 | **0.973** | **494.92** | **0.00** | **0.66** | **NULL** | **NA** | **-0.192** | **0.500** |
| 3 | 0.895 | 496.92 | 2.00 | 0.24 | NULL | 0.009 | -0.193 | 0.499 |
| 1 | 0.770 | 498.78 | 3.86 | 0.10 | 0.047 | 0.009 | -0.194 | 0.502 |
| 4 | 0.004 | 508.06 | 13.14 | 0.00 | NULL | 0.031 | -0.167 | NULL |
| 5 | 0.001 | 509.92 | 15.00 | 0.00 | 0.047 | 0.031 | -0.167 | 0.001 |
| Filter-Feeder |  |  |  |  |  |  |  |  |
| 5 | **0.467** | **496.24** | **0.00** | **0.36** | **0.254** | **-0.509** | **0.419** | **-0.129** |
| 4 | **0.272** | **496.61** | **0.37** | **0.30** | **NULL** | **-0.502** | **0.414** | **NULL** |
| 1 | **0.355** | **497.57** | **1.33** | **0.19** | **0.254** | **-0.530** | **0.400** | **-0.037** |
| 3 | **0.198** | **497.95** | **1.71** | **0.15** | **NULL** | **-0.528** | **0.399** | **0.098** |
| 2 | < 0.000 | 512.46 | 16.22 | 0.00 | NULL | NA | 0.337 | -0.021 |
| Shredder |  |  |  |  |  |  |  |  |
| 2 | **0.726** | **462.37** | **0.00** | **0.58** | **NULL** | **NA** | **0.021** | **0.652** |
| 3 | **0.759** | **463.61** | **1.24** | **0.31** | **NULL** | **0.104** | **0.033** | **0.635** |
| 1 | 0.459 | 465.60 | 3.23 | 0.11 | 0.160 | 0.104 | 0.033 | 0.652 |
| 4 | < 0.000 | 483.78 | 21.41 | 0.00 | NULL | 0.207 | 0.051 | NULL |
| 5 | < 0.000 | 485.78 | 23.41 | 0.00 | 0.160 | 0.207 | 0.051 | 0.033 |
| Predator |  |  |  |  |  |  |  |  |
| 1 | **0.664** | **504.15** | **0.00** | **0.39** | **0.129** | **0.136** | **0.239** | **0.434** |
| 2 | **0.228** | **504.29** | **0.14** | **0.36** | **NULL** | **NA** | **0.237** | **0.448** |
| 3 | **0.206** | **505.12** | **0.97** | **0.24** | **NULL** | **0.139** | **0.246** | **0.428** |
| 5 | 0.008 | 511.55 | 7.40 | 0.01 | 0.129 | 0.192 | 0.345 | 0.025 |
| 4 | 0.006 | 512.52 | 8.37 | 0.01 | NULL | 0.193 | 0.346 | NULL |

Note: All models in the re-analysis were compared for the full community and each functional feeding group. Models are ordered and formatted following Table 1.

## Drivers of β-Diversity: Ecological Context

Given the deviation between the results presented by Patrick and Brown (2018) and those presented here, I argue this could have been prevented by incorporating and applying proper ecological and statistical context. Regarding ecological context, Patrick and Brown (2018) neglected to present or provide summary data on community composition. Given the nature of the sampling design (i.e. first- to fourth-order streams), it is to be expected that collector-gatherers, shredders, and predators would comprise the majority of community abundance and biomass and play critical roles in food web structure and ecosystem functioning (Vannote et al. 1980, Rosi-Marshall and Wallace 2002). It would be more relevant to the ecology of the system to focus on the dominant taxa and concomitant functional feeding group subsets, in terms of relative abundance and biomass, rather than treating all functional feeding groups as equivalent. Based on my re-analysis of the complete model set, functional diversity only had a significantly-positive effect on driving β-diversity of collector-gatherers, and the magnitude of functional diversity on β-diversity (standardized path coefficient = 0.264) was roughly half of the magnitude of environmental heterogeneity (standardized path coefficient = 0.527); γ diversity had a significant role in driving β-diversity of shredders (standardized path coefficient = 0.652) and predators (standardized path coefficient range = 0.416-0.448).

## Drivers of β-Diversity: Statistical Context

Patrick and Brown (2018) also ignored important statistical context, partly derived from neglecting ecological context but also by conducting suspect analyses and subsequent presentation of results. First, I argue that applying the models to the whole community and each functional feeding group subset is a quasi-form of p-hacking. Rather than focusing on dominant taxa or groups relevant to the ecology of their study systems, Patrick and Brown (2018) had six model sets (i.e. the whole community and five functional feeding group subsets) in which to find a role for functional diversity driving β-diversity, enabling the identification of statistical significance without integrating ecological relevance. Moreover, only one out of five models would contradict their hypothesis, increasing the potential to find a statistically-significant but ecologically-contentious result. Second, I contend that the presence of a causal pathway between functional diversity and β-diversity is irrelevant because that does not indicate if the pathway is statistically significant, although this is purportedly how Patrick and Brown (2018) evaluated model results (see table 3). By not acknowledging if the pathway was significant, Patrick and Brown (2018) did not put the results into the appropriate context. Causal pathways could improve overall model fit, but it is fallacious to conflate improving model fit with having a significant effect on β-diversity, which is ostensibly the argument made by Patrick and Brown (2018). Finally, I claim the presentation of results is highly misleading. Patrick and Brown (2018) report the net effects of correlational pathways or pathways excluded from the respective model as 0.000 (see table 2). It is specious to report a net effect of 0 for a pathway that was not modeled, and the nature of correlational pathways in structural equation models precludes reporting direct or indirect effects mediated by correlational pathways (Grace 2006). Reporting net effects as null or not applicable, as done in this comment, would have been a better way to present the results as it accurately reflects model structures.

## Concluding Remarks

I contend that critical errors in the evaluation and subsequent interpretation of hypotheses led to spurious claims by overlooking fundamental ecological and statistical context. Although my alternative models were not consistently among the top models (table 1), I demonstrate that, as predicted, environmental heterogeneity and γ-diversity had consistently stronger and significant effects on β-diversity compared to effects mediated by functional diversity. Importantly, I show that results presented by Patrick and Brown (2018) are not reproducible, and this lack of reproducibility is concerning because it is avoidable. Patrick and Brown (2018) provided data and used open-access software for analyses, but analytical scripts were not archived alongside the data. Additionally, the supplementary data were partial; no raw environmental or community data were provided, limiting critical evaluation by reviewers and readers. Given the growing call for a culture shift with regards to open access, data archiving, and reproducibility in ecology and evolution (Hampton et al. 2015, Roche et al. 2015), publishing full data and analytical scripts should be done whenever possible to preclude the presentation of irreproducible results. In summary, I discourage the use of results presented by Patrick and Brown (2018) because further analyses demonstrate the results are incorrect. Although functional diversity is important for linking community and ecosystem ecology (McGill et al. 2006, Petchey and Gaston 2006), evidence from Patrick and Brown (2018) suggests functional diversity does not generate β-diversity. Instead, environmental variation and diversity within the regional species pool are the primary drivers underlying β-diversity.

## Appendix

Structural equation models (SEMs) were constructed to evaluate the causal pathways through which environmental variation, functional diversity, and γ-diversity structure β-diversity. Structural equation models are tests of fit to the data, and model fit was assessed by comparing expected and observed covariance between predictor and response variables using chi-square tests (Grace 2006). Structural equation models were considered consistent with the data when expected and observed covariance was not significantly different (i.e. p > 0.05). Hypothesized SEMs were then compared using Akaike’s information criterion (AIC, Burnham and Anderson 2002), ΔAIC (change in AIC value, Burnham and Anderson 2002), and AIC weight (relative support for the model, Burnham and Anderson 2002). Model selection occurred in two distinct phases. First, SEMs had to have good fit to the data (p > 0.05) and be considered statistically equivalent (ΔAIC < 2.00). Second, SEMs with comparable AIC weights were selected as the top models within the model set and then included in model interpretation and discussion. All SEMs were estimated by maximum likelihood. Path coefficients, which show the direction and magnitude of the causal relationship between variables, were standardized to allow for the comparison of relationship strengths within each SEM (Grace 2006). All variables were scaled (i.e. centered on the mean and then divided by the standard deviation) prior to analysis. Scaling was done on the full dataset and then on each functional feeding group subset individually. Statistical analyses were conducted using R (version 3.4.3, R Core Team 2017) using the lavaan (Rosseel 2012) and AICcmodavg (Mazerolle 2017) packages; R code for the re-analyses is deposited at figshare (10.6084/m9.figshare.6163841).

